# DNA looping mediates cooperative transcription activation

**DOI:** 10.1101/2022.10.10.511513

**Authors:** Shu-Jing Han, Yong-Liang Jiang, Lin-Lin You, Li-Qiang Shen, Feng Yang, Ning Cui, Wen-Wen Kong, Hui Sun, Ke Zhou, Hui-Chao Meng, Zhi-Peng Chen, Yuxing Chen, Yu Zhang, Cong-Zhao Zhou

## Abstract

Transcription factors respond to multi-level stimuli and co-occupy promoter regions of target genes to activate RNA polymerase (RNAP) in a cooperative manner. To decipher the molecular mechanism, here we report two cryo-electron microscopy structures of *Anabaena* transcription activation complexes (TACs): NtcA-TAC composed of RNAP holoenzyme, promoter and a global activator NtcA, and NtcA-NtcB-TAC comprising an extra context-specific regulator NtcB. Structural analysis showed that NtcA binding makes the promoter DNA bend by ∼50º, which facilitates RNAP to contact NtcB at the distal upstream NtcB box. The sequential binding of NtcA and NtcB induces looping back of promoter DNA towards RNAP, enabling the assembly of a fully activated TAC bound with two activators. Together with biochemical assays, we propose a ‘DNA looping’ mechanism of cooperative transcription activation in bacteria.

---

Multiple transcription factors act in concert via co-occupying the promoter regions and respond to different input signals to precisely control the transcription output of downstream genes in both prokaryotes and eukaryotes (*1-5*). In bacteria, one global regulator and one context-specific regulator usually function as a pair to activate gene transcription in a cooperative manner (*1, 3, 4*). Despite structures of TACs containing either class I or class II transcription activators have been reported (*6-8*), the underlying mechanism of cooperative transcription activation remains unclear due to the lack of structural information for a TAC comprising two activators.

Carbon/nitrogen (C/N) balance control is critical to maintain the cellular homeostasis for all organisms (*9*). As an ancient photoautotrophic bacterium, cyanobacteria have evolved a finely regulated signal transduction network to keep pace of nitrogen assimilation with CO_2_ fixation (*9-12*). The global regulator NtcA in the catabolite activator protein (CAP) family plays a central role in maintaining the C/N balance through sensing the central metabolite 2-oxoglutarate (2-OG), which tightly controls the expression of more than 120 genes that stimulate nitrogen metabolism in *Anabaena* sp. PCC 7120 (*Anabaena* for short) (*13-17*). Cyanobacteria mainly utilize the nitrate assimilation pathway encoded by *nirA-nrtABCD-narB* operon (termed *nirA* operon for short) to import and reduce nitrate (*10, 11, 18*). The expression of *nirA* operon is maintained at basal level by the global regulator NtcA but significantly boosted by the context-specific activator NtcB (a LysR-type transcriptional regulator; LTTR) when the nitrogen source is switched from ammonium to nitrate (*19-23*). Therefore, the regulation of *nirA* operon by NtcA and NtcB serves as an excellent model for studying cooperative transcription activation in bacteria.

We first verified that NtcA and NtcB activate transcription of the promoter of *nirA* operon (P*nirA*) in a cooperative manner. The *in vitro* transcription activity assays showed that 2-OG-bound NtcA increases the transcription activity of RNAP on P*nirA* (fig. S1A); and by contrast, NtcB alone has no effect on P*nirA* transcription (fig. S1B). However, NtcB substantially augments P*nirA* transcription in the presence of 2-OG-bound NtcA (fig. S1C). Our results showed that the transcription activation activity of NtcB relies on NtcA, confirming the cooperative model of action by NtcA and NtcB (*10, 19-22*).

To understand how NtcA activates transcription, we reconstituted the NtcA-TAC using *Anabaena* RNAP, σ^A^, 2-OG-bound NtcA, and the 125-bp P*nirA-*derived DNA comprising the consensus binding motifs of NtcA (-48 to -35 respective to the transcription start site, centered at -41.5) and NtcB (-100 to -64, centered at -82), also termed NtcA and NtcB boxes (Fig. 1A and fig. S1D). The cryo-EM map of NtcA-TAC reconstructed at 3.6 Å (fig. S2) shows unambiguous signals for 66-bp dsDNA (-53 to +13) with a partially resolved transcription bubble, RNAP and NtcA dimer (Fig. 1, B and C, and fig. S3A). The RNAP and the core promoter region (-36 to +13) adopt the similar conformation and make essentially the same interactions with each other as observed in previously reported open conformation of RNAP-promoter DNA (RPo) structure (fig. S3A) (*24*).

**Fig. 1.**
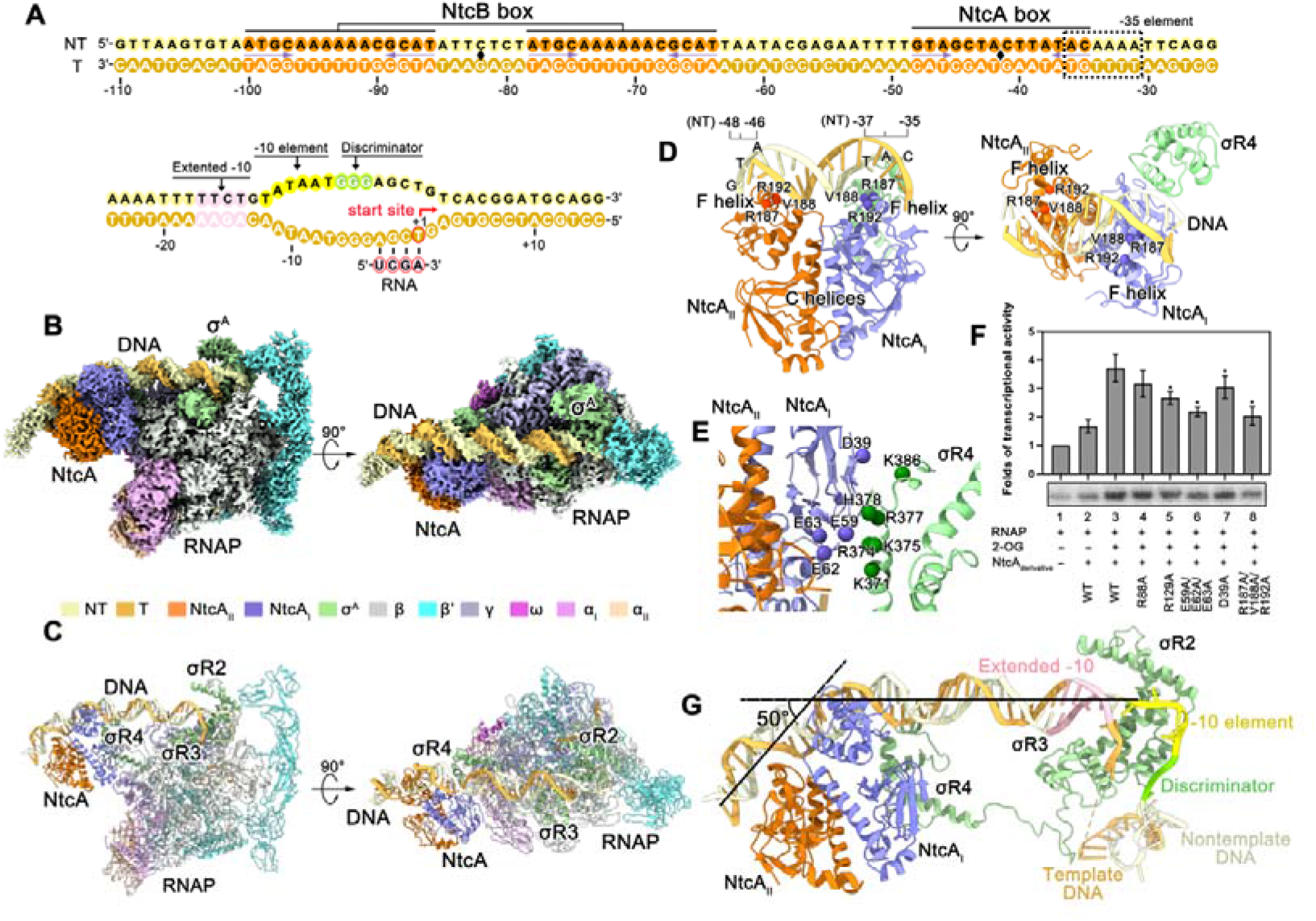
The structure of NtcA-TAC. **(A)** The nucleic-acid scaffold used for cryo-EM structure determination of NtcA-TAC and NtcA-NtcB-TAC. The NtcA and NtcB binding sites are highlighted and labeled with NtcA box and NtcB box, respectively. The palindromic sequences are underlined by purple arrows. The -35 element is highlighted in a dashed box. The **(B)** cryo-EM map and **(C)** the structure model of NtcA-TAC presented in two view orientations. The RNAP subunits, NtcA dimer and the promoter are colored differently as shown in the color key. The σR2, σR3, and σR4 represent the region 2, region 3 and region 4 of σ^A^, respectively. **(D)** NtcA and σR4 recognize partially overlapped NtcA box and the -35 element, respectively. NtcA dimer and σR4 are represented as cartoon. Cα atoms of the key interaction residues are represented as spheres. **(E)** Interactions between NtcA and σR4. Cα atoms of the interface residues are shows as spheres. **(F)** *In vitro* transcription activation results using a modified P*nirA* as template DNA and WT/mutant NtcA. Data are presented as mean ± S.E.M., n=3 biologically independent experiments. The lower panel shows the representative gel image. *p<0.05 indicates significant difference compared to lane 3. **(G)** The binding of NtcA dimer induces a 50° bending of the promoter DNA at the - 35 element.

NtcA functions as a class II activator on the P*nirA* promoter. The proximal protomer (NtcA_I_) of NtcA dimer makes interactions with the region 4 of σ^A^ (σR4) through one surface patch of NtcA that corresponds to the activating region 3 of *E. coli* CAP (Fig. 1, D and E, and fig. S3, B and C) (*25, 26*). The interactions involve residues Asp39, Glu59, Glu62, and Glu63 of NtcA and residues Lys371, Arg374, Lys375, Arg377, His378, and Lys386 of σR4 (Fig. 1E and fig. S3C). Mutating these interface residues on NtcA substantially impaired its transcription activation activity, indicating an important role of NtcA/σR4 interaction on the transcription activation activity of NtcA (Fig. 1F). Intriguingly, different from typical class II activators (*8, 26, 27*), NtcA makes interaction with neither the N-terminal (αNTD) nor C-terminal domain (αCTD) of α subunit of RNAP.

σR4 and NtcA recognize the -35 element and NtcA box, respectively, which are partially overlapped (Fig. 1, A and D, and fig. S3B). The σR4 loosely contacts the -35 element; and the helix-turn-helix (HTH) module of σR4 doesn’t reach the major groove as deeply as it does in the transcription initiation complex (RPitc) structure in *Synechocystis* sp. PCC 6803 (fig. S3D) (*24*), probably due to change of the dsDNA trajectory around the -35 element caused by the NtcA-DNA interactions, explaining a degenerated -35 sequence of P*nirA* promoter (Fig. 1A and fig. S1D). The degenerate - 35 motif and the weaken recognition by σR4 have also been observed in other class II TAC structures (*6, 28*). The DNA-binding domains (NtcA-DBDs) of NtcA dimer recognize the palindromic motif of the 14-bp NtcA box (^−48^GTA-*N*_*8*_-TAC^−35^) centered at -41.5. The ‘F helix’ and the side wing of the winged HTH module of NtcA-DBD insert into the major and minor grooves of the half palindromic NtcA box, respectively (Fig. 1D). Residues Arg187, Val188 and Arg192 of NtcA-DBD make direct interactions with DNA (fig. S3E). Alanine substitution of these residues not only greatly reduced the binding affinity of NtcA towards the promoter DNA (fig. S3F), but also lowered its transcription activation activity (Fig. 1F). Binding of NtcA dimer to the NtcA box enforces a ∼50º bending of DNA towards RNAP (Fig. 1G). This bending changes the trajectory of upstream dsDNA, shortens the spatial distance between RNAP and upstream DNA, and eventually makes RNAP accessible to the transcription factor that binds to the distal upstream DNA.

To understand how NtcB cooperates with NtcA to activate transcription, we determined the structure of a TAC comprising both NtcA and NtcB (NtcA-NtcB-TAC). The gel shift assays showed that sequential addition of NtcA and NtcB gradually slowed the migration of RNAP-promoter complex, suggesting formation of a larger complex (fig. S4). The reconstituted complex of NtcA-NtcB-TAC that comprises *Anabaena* RNAP, σ^A^, 2-OG-bound NtcA, NtcB, and the 125-bp P*nirA-*derived promoter DNA was subsequently applied to cryo-EM single particle analysis. The overall 4.5 Å cryo-EM map of NtcA-NtcB-TAC (fig. S5) shows unambiguous signals for RNAP, σ^A^, the NtcA dimer, the NtcB tetramer, ∼112-bp promoter DNA with a partially resolved transcription bubble (Fig. 2, A and B). The structural model of NtcA-NtcB-TAC was built by sequentially docking NtcA-TAC, the 2.4 Å crystal structure of the dimeric effector-binding domain of NtcB (NtcB-EBD) determined in this study (fig. S6, A and B; table S2), a predicted model of the DNA-binding domain of NtcB (NtcB-DBD) into the cryo-EM map. The structure shows that RNAP-core promoter interactions in the NtcA-NtcB-TAC remains unchanged compared to the structure of NtcA-TAC, suggesting that binding of NtcB does not alter RPo conformation (fig. S7A). The interactions of NtcA with RNAP and DNA are essentially the same in both NtcA-TAC and NtcA-NtcB-TAC structures, suggesting that NtcB binding doesn’t affect the binding mode of NtcA (fig. S7B).

**Fig. 2.**
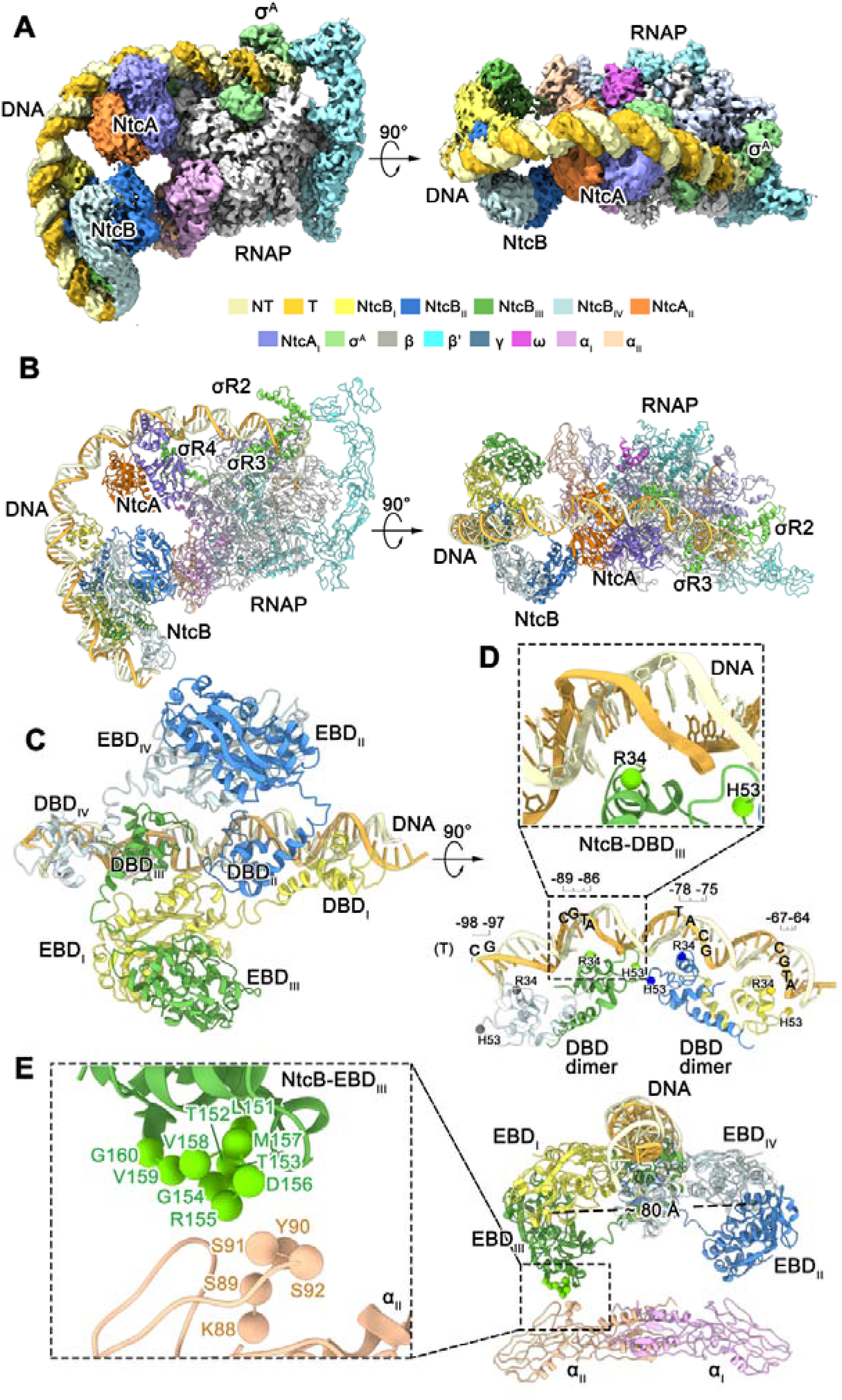
The structure of NtcA-NtcB-TAC. The **(A)** cryo-EM map and the **(B)** structure model of NtcA-NtcB-TAC are presented in two view orientations. The RNAP subunits, NtcA and NtcB protomers are colored as shown in the color key. **(C)** The NtcB tetramer binds to the NtcB box. Two NtcB-EBD dimers (NtcB-EBD_I_/NtcB-EBD_III_ and NtcB-EBD_II_/NtcB-EBD_IV_) are flanked at different sides of the promoter DNA. Two NtcB-DBD_III_/NtcB-DBD_IV_ and NtcB-DBD_I_/NtcB-DBD_II_ dimers are lined up along the promoter DNA. **(D)** The interactions between NtcB-DBDs and NtcB box with a zoom-in view shown as an inlet. The interface residues are shown as spheres. The positions of the conserved recognition motifs are highlighted. **(E)** The interactions between NtcB-EBD and RNAP-αNTD with a zoom-in view shown as an inlet. The potential interface residues are shown as spheres.

In NtcA-NtcB-TAC structure, NtcA and NtcB respectively interact with their boxes and slightly contact each other (Fig. 2, A and B). Beyond the interactions between RNAP holoenzyme and the core region of promoter, NtcA binds to σR4 and NtcA box, whereas NtcB makes interactions with RNAP-αNTD and NtcB box (Fig. 1 and Fig. 2). The electrophoretic mobility shift assays show that both NtcA and NtcB bind to P*nirA* in a sequence-specific manner (fig. S8, A and B). Moreover, mutating either NtcA or NtcB box did not affect DNA binding affinity of the other transcription factor (fig. S8, C and D). Our results suggest that NtcA and NtcB respectively bind to their boxes in an independent manner, different from the mutually dependent DNA binding mode of many eukaryotic transcription activators (*29-31*).

The LTTR-family regulator NtcB adopts a tetrameric structure that consists of two compact (NtcB_II_ and NtcB_III_) and two extended subunits (NtcB_I_ and NtcB_IV_), respectively (Fig. 2C). Two NtcB-EBD dimers (NtcB-EBD_I_/NtcB-EBD_III_ and NtcB-EBD_II_/NtcB-EBD_IV_) are flanked at the two different sides of promoter DNA and separated ∼80 Å in distance from each other, while two NtcB-DBD dimers (NtcB-DBD_III_/NtcB-DBD_IV_ and NtcB-DBD_I_/NtcB-DBD_II_) are lined up along the promoter DNA (Fig. 2, C and D). Generally, the NtcB tetramer adopts an expanded conformation that enables the binding to both the promoter DNA and RNAP via DBDs and EBDs, respectively.

The NtcA-NtcB-TAC structure reveals how NtcB interacts with DNA (Fig. 2, C and D). Similar to typical LTTRs (*23, 32*), each of two NtcB-DBD dimers is organized into a head-to-head manner to recognize the NtcB box (^−100^ATGC-*N*_*7*_-GCAT-*N*_*7*_-ATGC-*N*_*7*_-GCAT^−64^) that comprises two palindromic motifs interspaced by 7 bp (Fig. 1A and Fig. 2, C and D). The NtcB-DBD_III_/NtcB-DBD_IV_ dimer recognizes the distal palindromic box, while the NtcB-DBD_I_/NtcB-DBD_II_ dimer interacts with the proximal box (Fig. 2, C and D). Further structural analysis showed that residues Arg34 and His53 of NtcB-DBD, which are conserved in LTTRs (*23, 33*), directly interact with DNA (Fig. 2D and fig. S9A). Alanine substitution of these residues greatly reduced the binding affinity of NtcB towards the promoter (fig. S9B) and decreased its transcription activation activity (fig. S9C), confirming the essential roles of these residues.

The NtcA-NtcB-TAC structure also reveals how NtcB interacts with RNAP. Compared to the previously reported structures of LTTRs (*23, 34*), the two EBD dimers of NtcB are distantly separated whereas the two DBD dimers closely approach to each other (Fig. 2, D and E). The two separated EBD dimers make interactions with the two RNAP-α subunits via NtcB-EBD_II_ and NtcB-EBD_III_, respectively (Fig. 2E and fig. S9D). In the NtcB-EBD_I_/NtcB-EBD_III_ dimer, a small surface patch (residues Leu151-Gly160) of EBD_III_ approaches to the N-terminal domain of RNAP-α_II_ subunit (RNAP-αNTD_II_) and makes interactions most likely with a loop region (residues Lys88-Ser92) of RNAP-αNTD_II_ (Fig. 2E and fig. S9D). Similar interaction networks are also observed for the NtcB-EBD_II_/NtcB-EBD_IV_ dimer and RNAP-αNTD_I_ (Fig. 2E and fig. S9D). The interactions between RNAP-αNTD and NtcB-EBD are critical, as deletion of the interacting patch (residues Leu151-Gly160) of NtcB substantially impaired the cooperative transcription activation activity (fig. S9C).

The binding of NtcA and NtcB to the promoter respectively enforces a DNA bending of ∼50º and ∼70º at their binding boxes (fig. 3A). Moreover, NtcB binding to RNAP triggers an extra bending of ∼20º between NtcA and NtcB boxes. Therefore, the sequential binding of NtcA and NtcB induces a total DNA bending of ∼140°, resulting in looping back of the upstream promoter DNA in the NtcA-NtcB-TAC structure (Fig. 3A). This 140° bending enables the concomitant binding of NtcA and NtcB with both RNAP and promoter DNA. The 15-bp spacer between the binding boxes of NtcA and NtcB makes it possible that the NtcB tetramer and NtcA dimer bind to the promoter DNA on the same side that faces RNAP (Fig. 1A, 3A, and fig. S1D). Indeed, either extending or shortening the spacer by 5 bp completely abolished the cooperative activation activity of NtcB (fig. S10).

**Fig. 3.**
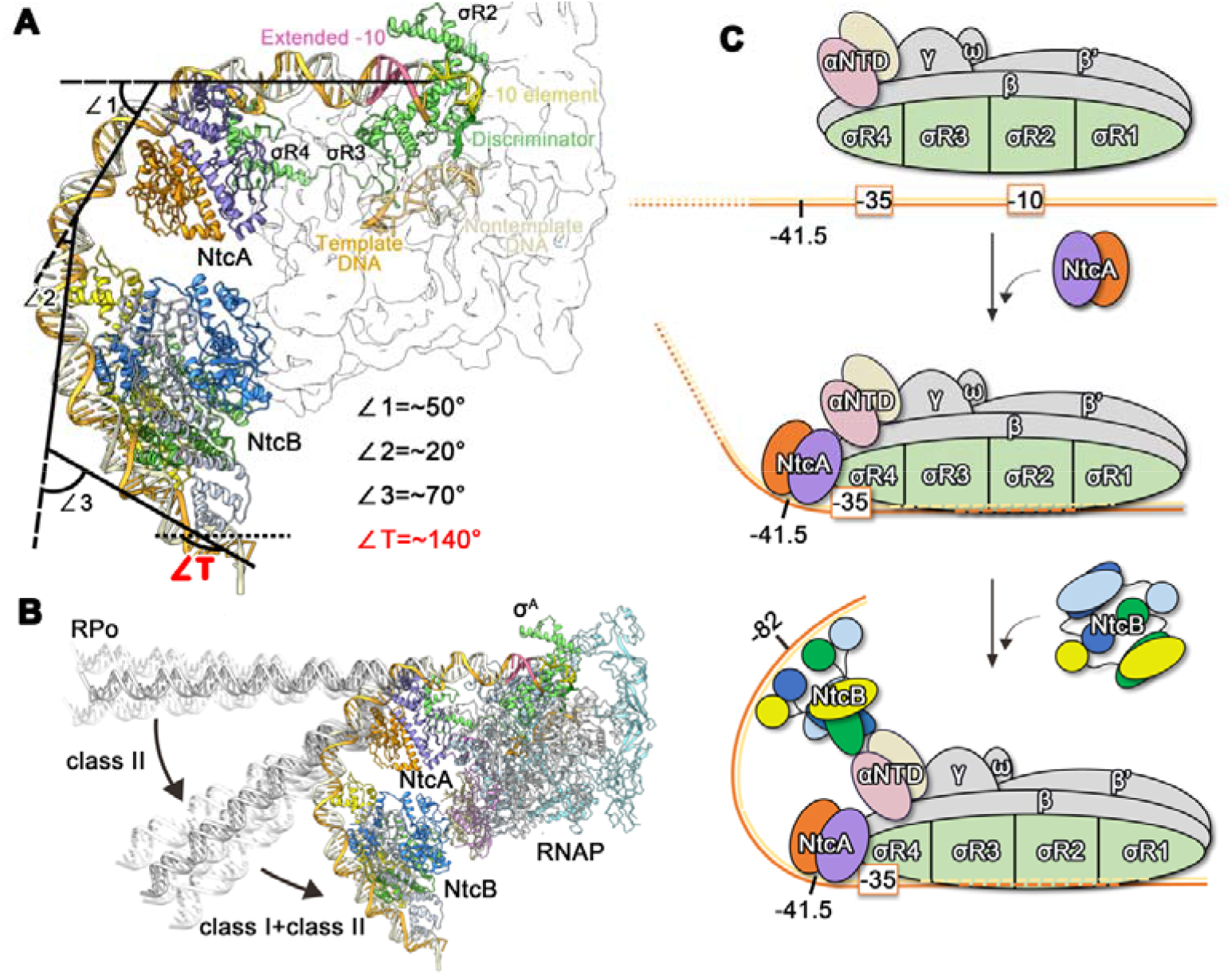
The ‘DNA looping’ model of cooperative transcription activation by NtcA and NtcB. **(A)** The upstream DNA is curved at positions -41.5 (∼50°), -61 (∼20°), -82 (∼70°), resulting in a total bending (✉T) of ∼140°. The promoter DNA, NtcA and NtcB protomers are colored as in Fig. 2. **(B)** Comparison of the upstream DNA trajectory of RPo, NtcA-TAC, and NtcA-NtcB-TAC. The bending of the promoter DNA is sequentially induced by the class II activator NtcA and the class I activator NtcB. The modeled promoter DNAs in flexible motion are colored in white. **(C)** The proposed ‘DNA looping’ model of cooperative transcription activation by NtcA and NtcB. The interactions among NtcA, RNAP, and promoter DNA induce a DNA bending by ∼50º, which enables RNAP to interact with NtcB located to the distal upstream promoter region. The sequential binding of NtcA and NtcB induces looping back of the promoter DNA towards RNAP, eventually enabling the assembly of a fully activated TAC bound with two activators.

These structural and biochemical analysis enabled us to propose a “DNA looping” mechanism underlying the cooperative activation by NtcA and NtcB, in which NtcB-RNAP interaction is enabled by the proximal promoter bending induced by NtcA-RNAP interaction (Fig. 3, B and C). The NtcB box is located at the distal upstream of the core promoter region, and thereby the tripartite interaction among NtcB, the promoter DNA, and RNAP-αNTD is energetically costly as it requires a DNA curving of ∼140º. Our structures show that NtcA bends the upstream DNA around its binding box and changes the trajectory of the upstream promoter towards RNAP (Fig. 3B). The NtcA-induced DNA bending significantly decreases the spatial distance between NtcB and RNAP-αNTD, making the interactions of which become possible. Moreover, it lowers the energetic barrier of DNA curving required for NtcB-RNAP interaction (Fig. 3, B and C). In the NtcA-NtcB-TAC structure, the extensive interactions among the activators, RNAP and promoter DNA are expected to stabilize the RNAP-DNA complex and facilitate subsequent promoter unwinding processes as in other class I and II bacterial transcription activation complexes (*6, 7, 35, 36*).

In summary, our findings reveal that cooperative activation of σ^70^−dependent transcription in bacteria involves a ‘DNA looping’ mechanism, by which the establishment of interactions between RNAP-σ^70^ holoenzyme and a class I activator bound at a distal upstream promoter region requires DNA looping by a class II activator bound at the proximal upstream promoter region. The ‘DNA looping’ model of cooperative activation has long been hypothesized in σ^54^−dependent transcription, in which the action of enhancer binding protein requires DNA looping by bacterial histone-like proteins (*1, 37*). This model has also long been hypothesized in eukaryotic Pol II transcription, in which the enhancer and promoter are brought together by DNA-looping forming proteins, topologically associating domains, as well as condensates (*2, 29, 38-40*). Our findings suggest a general means for transcription cooperative activation shared in bacteria and eukaryotes. Moreover, our work provides a detailed structural example on the ‘DNA looping’-mediated cooperative transcription activation.

## Acknowledgments

We thank the staff at the Shanghai Synchrotron Radiation Facility (SSRF) for X-ray diffraction data collection of NtcB-EBD; Pei-Ping Tang and Yong-Xiang Gao at the Cryo-EM Center at University of Science and Technology of China for cryo-EM data acquisition. This work is supported by the Strategic Priority Research Program of the Chinese Academy of Sciences (http://www.cas.cn; grant numbers XDB37020301 and XDA24020302), the National Natural Science Foundation of China (http://www.nsfc.gov.cn; grant number 32171198), National Key Research and Development Program of China (grant number 2018YFA090070), and Anhui Provincial Natural Science Foundation (http://kjt.ah.gov.cn; grant number 2108085J14). Y.-L.J. thanks the Youth Innovation Promotion Association of Chinese Academy of Sciences for their support (Membership No. 2020452).

## Author Contributions

C.-Z.Z., Y.Z., Y.-L.J. and Y.C. conceived, designed, and supervised the project. C.-Z.Z., Y.Z., Y.-L.J., Y.C. and S.-J.H. analyzed data and wrote the manuscript. S.-J.H., L.-Q.S., H.S., H.-C.M., K.Z., and N.C. performed the molecular cloning, protein expression and purification. S.-J.H., L.-L.Y. and W.-W.K. conducted the cryo-EM sample preparation and data acquisition. S.-J.H., Y.-L.J. and F.Y. performed cryo-EM data processing and model building. S.-J.H. performed the biochemical assays, protein crystallization and optimization. Y.-L.J. and Z.-P.C conducted the X-ray data collection, Y.-L.J. performed crystal structure determination and model building. All authors discussed the data and read the manuscript.

## Data availability

The crystal structure of NtcB-EBD has been deposited at Protein Data Bank (PDB) under the accession code of 8H3Z. The cryo-EM maps and coordinates of NtcA-TAC and NtcA-NtcB-TAC have been deposited at the Electron Microscopy Data Bank (EMDB) (EMD-34476 for NtcA-TAC and EMD-34475 for NtcA-NtcB-TAC) and PDB (8H40 for NtcA-TAC and 8H3V for NtcA-NtcB-TAC). The supplementary map of NtcA-NtcB-TAC focusing on NtcA and NtcB has been deposited at EMDB under accession code of EMD-34477.

## References

1. D. F. Browning, S. J. Busby, Local and global regulation of transcription initiation in bacteria. Nat Rev Microbiol 14, 638–650 (2016).

2. P. Cramer, Organization and regulation of gene transcription. Nature 573, 45–54 (2019).

3. D. F. Browning, S. J. Busby, The regulation of bacterial transcription initiation. Nat Rev Microbiol 2, 57–65 (2004).

4. D. J. Lee, S. D. Minchin, S. J. Busby, Activating transcription in bacteria. Annu Rev Microbiol 66, 125–152 (2012).

5. S. J. W. Busby, Transcription activation in bacteria: ancient and modern. Microbiology 165, 386–395 (2019).

6. Y. Feng, Y. Zhang, R. H. Ebright, Structural basis of transcription activation. Science 352, 1330–1333 (2016).

7. B. Liu, C. Hong, R. K. Huang, Z. Yu, T. A. Steitz, Structural basis of bacterial transcription activation. Science 358, 947–951 (2017).

8. W. Shi, Y. Jiang, Y. Deng, Z. Dong, B. Liu, Visualization of two architectures in class-II CAP-dependent transcription activation. PLoS Biol 18 (2020).

9. K. Forchhammer, K. A. Selim, Carbon/nitrogen homeostasis control in cyanobacteria. FEMS Microbiol Rev 44, 33–53 (2020).

10. Y. Ohashi et al., Regulation of nitrate assimilation in cyanobacteria. J Exp Bot 62, 1411–1424 (2011).

11. E. Flores, J. E. Frias, L. M. Rubio, A. Herrero, Photosynthetic nitrate assimilation in cyanobacteria. Photosynth Res 83, 117–133 (2005).

12. E. Flores, A. Herrero, Nitrogen assimilation and nitrogen control in cyanobacteria. Biochem Soc Trans 33, 164–167 (2005).

13. M. X. Zhao et al., Structural basis for the allosteric control of the global transcription factor NtcA by the nitrogen starvation signal 2-oxoglutarate. Proc Natl Acad Sci U S A 107, 12487–12492 (2010).

14. J. L. Llacer et al., Structural basis for the regulation of NtcA-dependent transcription by proteins PipX and PII. Proc Natl Acad Sci U S A 107, 15397–15402 (2010).

15. S. Picossi, E. Flores, A. Herrero, ChIP analysis unravels an exceptionally wide distribution of DNA binding sites for the NtcA transcription factor in a heterocyst-forming cyanobacterium. BMC Genomics 15, (2014).

16. M. A. Dominguez-Martin et al., Differential NtcA Responsiveness to 2-Oxoglutarate Underlies the Diversity of C/N Balance Regulation in Prochlorococcus. Front Microbiol 8, (2017).

17. J. Giner-Lamia et al., Identification of the direct regulon of NtcA during early acclimation to nitrogen starvation in the cyanobacterium Synechocystis sp. PCC 6803. Nucleic Acids Res 45, 11800–11820 (2017).

18. S. Imamura et al., Nitrate assimilatory genes and their transcriptional regulation in a unicellular red alga Cyanidioschyzon merolae: genetic evidence for nitrite reduction by a sulfite reductase-like enzyme. Plant Cell Physiol 51, 707–717 (2010).

19. K. V. Lopatovskaia, A. V. Seliverstov, V. A. Liubetskii, NtcA- and NtcB-regulons in cyanobacteria and Rhodophyta chloroplasts. Mol Biol 45, 570–574 (2011).

20. J. E. Frias, E. Flores, A. Herrero, Activation of the Anabaena nir operon promoter requires both NtcA (CAP family) and NtcB (LysR family) transcription factors. Mol Microbiol 38, 613–625 (2000).

21. M. Aichi, T. Omata, Involvement of NtcB, a LysR family transcription factor, in nitrite activation of the nitrate assimilation operon in the cyanobacterium Synechococcus sp. strain PCC 7942. J Bacteriol 179, 4671–4675 (1997).

22. M. Aichi, N. Takatani, T. Omata, Role of NtcB in activation of nitrate assimilation genes in the cyanobacterium Synechocystis sp. strain PCC 6803. J Bacteriol 183, 5840–5847 (2001).

23. S. E. Maddocks, P. C. F. Oyston, Structure and function of the LysR-type transcriptional regulator (LTTR) family proteins. Microbiology 154, 3609–3623 (2008).

24. L. Shen et al., https://biorxiv.org/cgi/content/short/2022.10.06.511230v1. (2022).

25. C. L. Lawson et al., Catabolite activator protein: DNA binding and transcription activation. Curr Opin Struct Biol 14, 10–20 (2004).

26. S. Busby, R. H. Ebright, Transcription activation by catabolite activator protein (CAP). J Mol Biol 293, 199–213 (1999).

27. Y. Feng, Y. Zhang, R.H. Ebright, Structural basis of transcription activation. Science 352 (2016).

28. G. Blanco, A. Canals, J. Bernues, M. Sola, M. Coll, The structure of a transcription activation subcomplex reveals how sigma(70) is recruited to PhoB promoters. EMBO J 30, 3776–3785 (2011).

29. Panigrahi, B. W. O’Malley, Mechanisms of enhancer action: the known and the unknown. Genome Biol 22, 108 (2021).

30. V. Haberle, A. Stark, Eukaryotic core promoters and the functional basis of transcription initiation. Nat Rev Mol Cell Biol 19, 621–637 (2018).

31. T. W. Sikorski, S. Buratowski, The basal initiation machinery: beyond the general transcription factors. Curr Opin Cell Biol 21, 344–351 (2009).

32. E. A. Giannopoulou et al., Crystal structure of the full-length LysR-type transcription regulator CbnR in complex with promoter DNA. FEBS J 288, 4560–4575 (2021).

33. M. Alanazi, E. L. Neidle, C. Momany, The DNA-binding domain of BenM reveals the structural basis for the recognition of a T-N11-A sequence motif by LysR-type transcriptional regulators. Acta Crystallogr D Biol Crystallogr 69, 1995–2007 (2013).

34. Y. L. Jiang et al., Coordinating carbon and nitrogen metabolic signaling through the cyanobacterial global repressor NdhR. Proc Natl Acad Sci U S A 115, 403–408 (2018).

35. Mazumder, A. N. Kapanidis, Recent Advances in Understanding sigma70-Dependent Transcription Initiation Mechanisms. J Mol Biol 431, 3947–3959 (2019).

36. M. Hao et al., Structures of Class I and Class II Transcription Complexes Reveal the Molecular Basis of RamA-Dependent Transcription Activation. Adv Sci (Weinh) 9, (2022).

37. E. Danson, M. Jovanovic, M. Buck, X. Zhang, Mechanisms of sigma(54)-Dependent Transcription Initiation and Regulation. J Mol Biol 431, 3960–3974 (2019).

38. S. Weintraub et al., YY1 Is a Structural Regulator of Enhancer-Promoter Loops. Cell 171, 1573–1588 (2017).

39. X. Chen et al., Structures of the human Mediator and Mediator-bound preinitiation complex. Science 372, (2021).

40. S. Schoenfelder, P. Fraser, Long-range enhancer-promoter contacts in gene expression control. Nat Rev Genet 20, 437–455 (2019).

