## Supplementary for "DNA looping mediates cooperative transcription activation"

**Materials and methods**

**Plasmid construction**

The plasmid information is summarized in table S1. The genes encoding α subunit (*rpoA*), β subunit (*rpoB*), γ subunit (*rpoC1*), β' subunit (*rpoC2*), ω subunit (*rpoZ*), σ factor (*sigA*, σ*^A^*), NtcA (*ntcA*), NtcB (*ntcB*) were amplified by PCR from *Anabaena* genomic DNA, and their mutants (NtcA^E59A/E62A/E63A^, NtcA^D39A^, NtcA^R187A/V188A/R192A^, NtcB^∆Leu151-Gly160^, NtcB^R34A/H53A^) were amplified by PCR from their wild-type plasmids, respectively. *Anabaena rpoA* and *rpoZ* were cloned into pCDFDuet^TM^-1 (Novagen); *rpoB*, *rpoC1,* and *rpoC2* were cloned into pETDuet (Novagen) using homologous recombination methods. *Anabaena* *ntcA*, *ntcB,* σ*^A^* and their derivatives were respectively cloned into the pET28a vector with an N-terminal His-tag.

pEASY/P*nirA* was reformed by pEASY-blunt vector (Transgen Biotech) The pEASY/P*nirA^AB+5^* and pEASY/P*nirA^AB-5^* were prepared using pEASY/P*nirA* as template through site-directed mutation.

**Protein expression and purification**

All proteins were expressed in *E. coli* strain BL21 (DE3) (Novagen). Protein expression was induced for 16 hours by addition of 0.4 mM isopropy β-D-1-thiogalactopyranoside (IPTG) at 18℃ when the optical density at 600 nm reaches 0.6 to 0.8. Cells were harvested by centrifugation (8,000 rpm, 3 min, 4℃). Cell pellets of NtcA, NtcB and their mutants were resuspended using the lysis buffer A and lysed by 30 min of sonication. After centrifugation at 12,000 rpm for 30 min, supernatants containing target proteins were loaded onto a Ni-NTA column (Cytiva) respectively equilibrated with lysis buffer A. Target proteins were eluted using lysis buffer A with 0.5 M imidazole and further loaded onto a Superdex 200 column (GE Healthcare) pre-equilibrated with lysis buffer A. Target protein fractions were eluted with buffer A, pooled, and concentrated to 10 mg mL^-1^. Cell pellet of NtcB-EBD was resuspended using the lysis buffer B and lysed by 30 min of sonication and purified by following above procedure.

*Anabaena* RNAP core enzyme was overexpressed in *E. coli* BL21 (DE3) cells carrying pETDuet/*rpoB-rpoC1-rpoC2* and pCDFDuet-1/*rpoA-rpoZ* and purified as described previously. *Anabaena* σ^A^ was overexpressed in *E. coli* BL21 (DE3) cells carrying pET28a/σ*^A^* and purified by a similar procedure for NtcA and NtcB.

**Crystallization and structure determination of NtcB-EBD**

Crystals of *Anabaena* NtcB-EBD were grown at 16°C by sitting drop vapor diffusion in a 2-µL drop. X-ray diffraction data were collected at 100 K at beamline 19U with a DECTRIS PILATUS 6M detector at the Shanghai Synchrotron Radiation Facility (SSRF). The diffraction data were integrated and scaled using XDS (*1*). The crystal structure of NtcB-EBD was determined by experimental phasing using PHENIX AutoSol (*2*). The initial model was built by AutoBuild, refined in REFMAC5 (*3*), and manually rebuilt interactively using the program Coot (*4*). The final model showed well geometry and were evaluated using MolProbity (http://molprobity.biochem.duke.edu). A list of the parameters of data collection, processing, structure determination and refinement is provided in table S2.

**Nucleic-acid scaffold**

Nucleic-acid scaffold for *Anabaena* NtcA-NtcB-TAC and NtcA-TAC was prepared as follows: nontemplate-strand DNA (NT-DNA, Sangon Biotech), template-strand DNA (T-DNA, Sangon Biotech) and oligoribonucleotide (5’-UCGA-3’; GenScript) in 42.4 μL annealing buffer with the ratio of 1.2:1:1.3 were heated for 5 min at 95℃ and cooled to 25℃ in 1℃ steps with 30 sec per step using a thermal cycler. The sequences of template and nontemplate DNAs are listed in table S1.

**Electrophoretic mobility shift assays (EMSA)**

Cy5-labeld DNA fragments used in EMSA assays for NtcB and NtcA binding were listed in table S1. The reaction mixture (10 μL) includes DNA fragment (0.2 μM, Sangon Biotech), WT/mutant NtcA (0, 0.5, 1, 2, 4 μM), or WT/mutant NtcB (0, 2, 4, 8, 16, 32 μM) in 20 mM Tris-HCl pH 8.0, 100 mM NaCl. The mixture was incubated for 20 min on ice. The solutions were separated by 5% Tris-boric acid (TBE) gel in the TBE buffer (89 mM Tris-boric pH 8.3, 2 mM EDTA) and analyzed by Tanon-2500 (Tanon Science & Technology).

***In vitro* transcription assays**

To measure the transcription activation activity of *Anabaena* NtcA, reaction mixtures (20 μL) containing RNAP-σ^A^ holoenzyme (25 nM; final concentration), 40 mM Tris-HCl, pH 8.0, 5 mM MgCl_2_, 75 mM NaCl, 12.5% (v/v) glycerol and 2.5 mM DTT, and P*nirA* (25 nM; final concentration; prepared by PCR using pEASY/P*nirA* as a template) were incubated for 10 min at 37℃. NtcA (0, 25, 50, 100, 200, 400, or 800 nM; final concentration) preincubated with α-ketoglutaric acid (2-OG, 1 µM) was added into the mixture followed by 10-min incubation at 37℃. The reactions were initiated by adding NTP mixture (100 μM [α-^32^P] UTP (0.04 Bq fmol^-1^), 100 μM of ATP, GTP and CTP; final concentration) and terminated after 10 min at 37℃ by addition of 5 μL of loading buffer (8 M urea, 20 mM EDTA, 0.025% xylene cyanol and 0.025% bromophenol blue). The samples were boiled for 2 min, and immediately cooled in ice for 5 min.

To measure the transcription activation activity by NtcB, reaction mixtures (20 μL) containing RNAP-σ^A^ holoenzyme (50 nM; final concentration), 40 mM Tris-HCl, pH 8.0, 5 mM MgCl_2_, 75 mM NaCl, 12.5% (v/v) glycerol, 2.5 mM DTT and P*nirA* or its derivatives (50 nM; final concentration; prepared by PCR using pEASY/P*nirA,* pEASY/P*nirA^AB+5^* and pEASY/P*nirA^AB-5^* as templates, respectively) were incubated for 10 min at 37℃. NtcB (0, 50, 100, 200, 400, 800, or 1600 nM; final concentration) and NtcA (200 nM; final concentraion) preincubated with 2-OG (2 µM; final concentration) were added into the reaction mixture followed by 10-min incubation at 37℃. The reactions were initiated by adding NTP mixture (100 μM [α-^32^P] UTP (0.04 Bq fmol^-1^), 100 μM of ATP, GTP and CTP; final concentration) and terminated after 10 min at 37℃ by addition of 5 μL of loading buffer (8 M urea, 20 mM EDTA, 0.025% xylene cyanol and 0.025% bromophenol blue). The samples were boiled for 2 min, and immediately cooled in ice for 5 min.

To measure the transcription activation activity by NtcA mutants, reaction mixtures (20 μL) containing RNAP-σ^A^ holoenzyme (25 nM; final concentration), 40 mM Tris-HCl, pH 8.0, 5 mM MgCl_2_, 75 mM NaCl, 12.5% (v/v) glycerol, 2.5 mM DTT and P*nirA* (25 nM; final concentration; prepared by PCR using pEASY/P*nirA* as template) were incubated for 10 min at 37℃. WT/mutant NtcA (100 nM; final concentration) preincubated with 2-OG (1 µM; final concentration) were added into the reaction mixture followed by 10-min incubation at 37℃. The reactions were initiated by adding NTP mixture (100 μM [α-^32^P] UTP (0.04 Bq fmol^-1^), 100 μM of ATP, GTP and CTP; final concentration) and terminated after 10 min at 37℃ by addition of 5 μL of loading buffer (8 M urea, 20 mM EDTA, 0.025% xylene cyanol and 0.025% bromophenol blue). The samples were boiled for 2 min, and immediately cooled in ice for 5 min.

To measure the transcription activation activity by NtcB mutants, reaction mixtures (20 μL) containing RNAP-σ^A^ holoenzyme (25 nM; final concentration), 40 mM Tris-HCl, pH 8.0, 5 mM MgCl_2_, 75 mM NaCl, 12.5% (v/v) glycerol, 2.5 mM DTT and P*nirA* or its derivatives (25 nM; final concentration; prepared by PCR using pEASY/P*nirA* as templates) were incubated for 10 min at 37℃. NtcA (50 nM; final concentraion) and WT/mutant NtcB (400 nM; final concentraion) preincubated with 2-OG (50 mM; final concentration) were added into the reaction mixture followed by 10-min incubation at 37℃. The reactions were initiated by adding NTP mixture (100 μM [α-^32^P] UTP (0.04 Bq fmol^-1^), 100 μM of ATP, GTP and CTP; final concentration) and terminated after 10 min at 37℃ by addition of 5 μL of loading buffer (8 M urea, 20 mM EDTA, 0.025% xylene cyanol and 0.025% bromophenol blue). The samples were boiled for 2 min, and immediately cooled in ice for 5 min.

RNA transcripts were separated in 15% (19:1 acrylamide/bisacrylamide) urea-polyacrylamide slab gels in 90 mM Tris-borate (pH 8.0) and 0.2 mM EDTA. The radiograph was obtained by storage-phosphor scanning (Typhoon; GE Healthcare) and the intensity was quantified by FUJIFILM Multi Gauge.

**Cryo-EM sample preparation and data collection**

The NtcA-TAC complex was reconstituted by mixing *Anabaena* RNAP-σ^A^ holoenzyme, NtcA, and nucleic-acid scaffold in a molar ratio of 1:4:1.3, while NtcA-NtcB-TAC was reconstituted mixing *Anabaena* RNAP-σ^A^ holoenzyme, NtcA, NtcB and nucleic-acid scaffold in a molar ratio of 1:4:10:1.3. Then 5 mM 2-OG was added to the reconstituted mixture and incubated for 2 hours on ice.

The NtcA-TAC and NtcA-NtcB-TAC complexes were further purified and cross-linked by Gradient Fixation (GraFix) (*5*). In brief, the solution was added to the top of a continuous density gradient (0-30% (v/v) glycerol and 0.05-0.02% paraformaldehyde) in buffer C. Then the mixture was centrifuged at 33,000 rpm for 16 hours at 4℃ using SW40Ti rotor (Beckman Coulter). The fractions containing target complex were collected and loaded on Superose 6 10/300 GL column (GE Healthcare) in buffer C to remove glycerol. The fractions of NtcA-TAC and NtcA-NtcB-TAC were pooled and concentrated to ~15 and ~17 mg mL^-1^, respectively.

The freshly prepared complexes were incubated with 3-([3-cholamidopropyl] dimethylammonio) −2-hydroxy-1-propanesulfonate (CHAPSO; Hampton Research, Inc) before grid preparation (*6*). Then, a drop of 3 μL was placed on a glow-discharged holey gold grid (UltrAuFoil, R 1.2/1.3 Au 400 mesh, Protochips), blotted with Vitrobot Mark IV (FEI) and plunge-frozen into liquid ethane with 100% chamber humidity at 4°C. The grids were blotted for 0.5 sec with a blot force of -1 and a wait time of 5 sec.

The NtcA-TAC and NtcA-NtcB-TAC cryo-EM datasets were collected using EPU on a 300-keV Titan Krios (FEI) equipped at the University of Science and Technology of China (USTC). For NtcA-TAC, a total of 2090 movies (32 frames, each 0.16 sec, total dose ~50 e Å^-2^) were recorded using a Gatan K3 Summit direct electron detector in super-resolution mode at a nominal magnification of ×81,000 at a pixel size of 0.535 Å and with a defocus range from -1.2 to -2.2 μm. For NtcA-NtcB-TAC, a total of 5047 movies (40 frames, each 0.16 sec, total dose ~50 e Å^-2^) were recorded using a Gatan K3 Summit direct electron detector in super-resolution mode at a nominal magnification of ×81,000 at a pixel size of 0.535 Å and with a defocus range from -1.2 to -2.2 μm.

**Cryo-EM data processing**

All of the movies were motion-corrected and dose-weighted using MotionCor2 (*7*) and were binned two-fold to yield a pixel size of 1.07 Å. The defocus values were estimated using CTFFIND4 (*8*).

For the dataset of NtcA-TAC, a total of 1,462 NtcA-TAC micrographs were processed for particle auto-picking using RELION after removing bad images by manual checking (*9*). Approximately 566,246 particles were boxed for 2D classification. All good classes containing 555,921 particles were selected for the first round 3D classification with four classes by C1 symmetry. 298,068 particles represented the complex containing RNAP and NtcA, whereas the rest of the three classes represented irrelevant junk particles. Approximately ~82.4% particles were selected for the second-round 3D classification and refinement, yielding a map of NtcA-TAC at 3.4 Å resolution after a PostProcess step as determined by Golden standard Fourier shell correlation using the 0.143 threshold. However, the cryo-EM map shows weak signal for the upstream portion of NtcA. To obtain a better map near NtcA region, we performed the third-round 3D classification without alignment and yielded 211,399 particles for the next masked 3D classification focusing on NtcA. ~21.4% particles were re-extracted to perform 3D refinement with solvent flattened FSCs, CTF refinement and Bayesian polishing, yielding a final map at resolution of 3.6 Å.

For the dataset of NtcA-NtcB-TAC, a total of 402,817 particles were selected after removing bad images and irrelevant junk particles for the first round 3D classification with four classes by C1 symmetry from 5047 images. Approximately ~16.2% particles represented the complex containing RNAP, NtcA and NtcB. The particles were imported into CryoSparc for the next 3D refinement and local refinement, yielding a final map at resolution of 4.5 Å. To obtain a better map at the region of NtcA and NtcB binding sites, 253,244 particles were selected to perform 3D classification focusing on the NtcA and NtcB binding regions without alignment. Totally 55,966 particles were selected for the final 3D refinement by imposing C1 symmetry, yielding a map at resolution of 7.6 Å.

**Model Building and Refinement**

Model building of NtcA-TAC was performed using Chimera (*10*) by manually fitting the *Synechocystis* sp. PCC 6803 RPitc (PDB: 8GZG) into the cryo-EM map. NtcA dimer was unambiguously docked into the map using the previously solved crystal structure of *Anabaena* sp. PCC 7120 NtcA (PDB: 3LA2) (*11*). Model building of the intact NtcA-NtcB-TAC was performed using Chimera by manually fitting the NtcA-TAC into the map. The structural model of full-length NtcB was constructed by a predicted model of NtcB-DBD and the crystal structure of NtcB-EBD (PDB: 8H3Z). The chimeric NtcB model was subsequently docked into the map in Chimera. The two structure models were adjusted in Coot (*4*), followed by the iterative positional and B-factor refinement in real space using PHENIX (*2*). The geometries of the final structures were evaluated using MolProbity (http://molprobity.biochem.duke.edu). A list of the parameters of cryo-EM data collection, processing, structure determination and refinement is provided in table S3.

References

1.	W. Kabsch, Xds. *Acta Crystallogr D Biol Crystallogr* **66**, 125-132 (2010).

2.	P. D. Adams *et al.*, PHENIX: building new software for automated crystallographic structure determination. *Acta Crystallogr D Biol Crystallogr* **58**, 1948-1954 (2002).

3.	G. N. Murshudov, A. A. Vagin, E. J. Dodson, Refinement of macromolecular structures by the maximum-likelihood method. *Acta Crystallogr D Biol Crystallogr* **53**, 240-255 (1997).

4.	P. Emsley, K. Cowtan, Coot: model-building tools for molecular graphics. *Acta Crystallogr D Biol Crystallogr* **60**, 2126-2132 (2004).

5.	H. Stark, GraFix: stabilization of fragile macromolecular complexes for single particle cryo-EM. *Methods Enzymol* **481**, 109-126 (2010).

6.	J. Chen, A. J. Noble, J. Y. Kang, S. A. Darst, Eliminating effects of particle adsorption to the air/water interface in single-particle cryo-electron microscopy: Bacterial RNA polymerase and CHAPSO. *J Struct Biol X* **1**, (2019).

7.	S. Q. Zheng *et al.*, MotionCor2: anisotropic correction of beam-induced motion for improved cryo-electron microscopy. *Nat Methods* **14**, 331-332 (2017).

8.	A. Rohou, N. Grigorieff, CTFFIND4: Fast and accurate defocus estimation from electron micrographs. *J Struct Biol* **192**, 216-221 (2015).

9.	S. H. Scheres, RELION: implementation of a Bayesian approach to cryo-EM structure determination. *J Struct Biol* **180**, 519-530 (2012).

10.	E. F. Pettersen *et al.*, UCSF Chimera–a visualization system for exploratory research and analysis. *J Comput Chem* **25**, 1605-1612 (2004).

11.	M. X. Zhao *et al.*, Structural basis for the allosteric control of the global transcription factor NtcA by the nitrogen starvation signal 2-oxoglutarate. *Proc Natl Acad Sci U S A* **107**, 12487-12492 (2010).

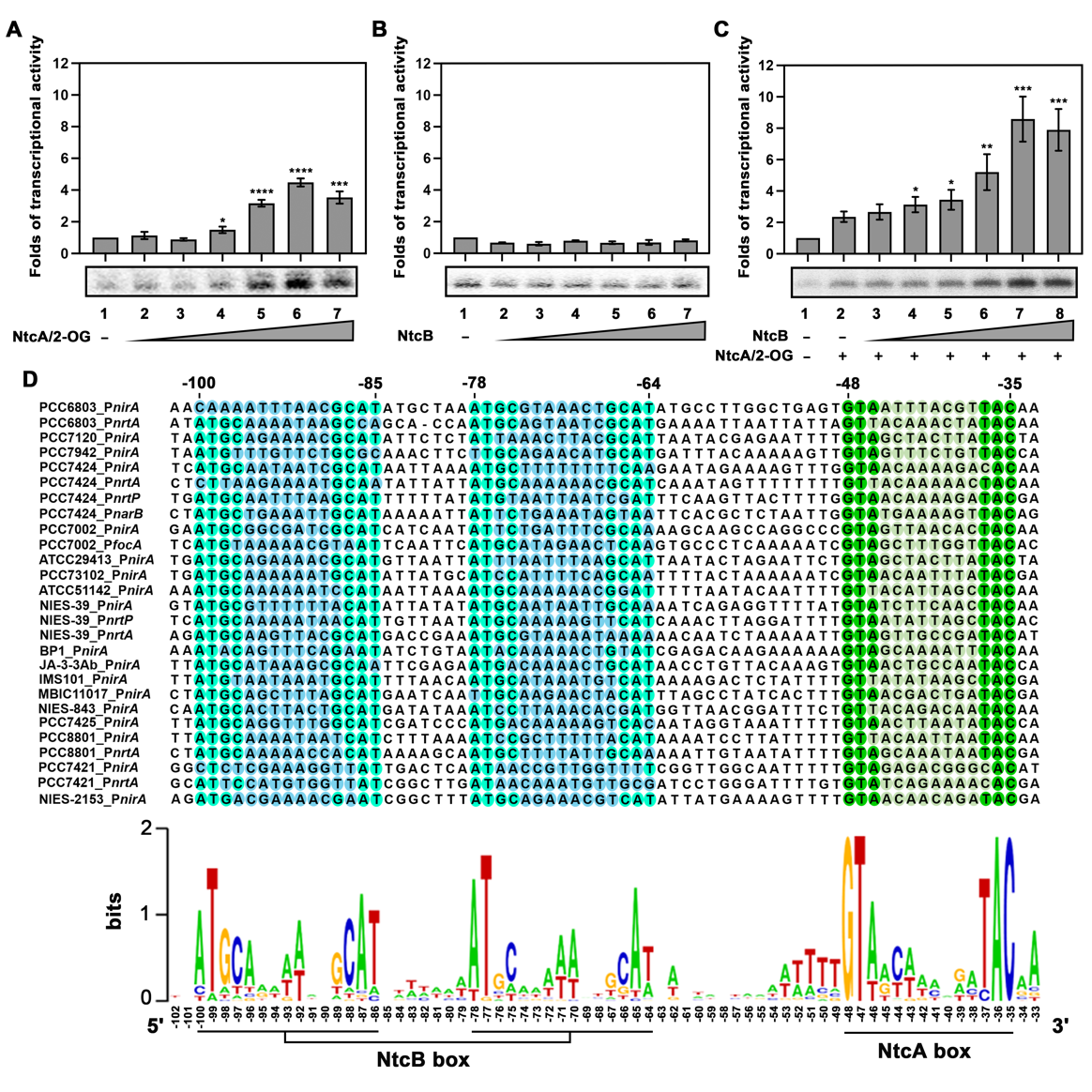

**fig. S1. The coordinated regulation of NtcA and NtcB on a P*nirA* derivative.** The transcriptional activity of *Anabaena* sp. PCC 7120 RNAP in the presence of (**A**) NtcA or (**B**) NtcB at increasing concentrations (0, 25, 50, 100, 200, 400, and 800 nM). (**C**) The transcriptional activity of RNAP preincubated with 100 nM NtcA, followed by adding NtcB at increasing concentrations (0, 50, 100, 200, 400, 800, and 1600 nM). Data are presented as mean ± S.E.M., n= 3 biologically independent experiments. The lower panel shows the representative gel image. *p<0.05, **p<0.01, ***p<0.001, and ****p<0.0001 indicate significant difference compared to lane 1. (**D**) The multiple-sequence alignment of the upstream promoter region of nitrate assimilation genes in β-cyanobacteria. The NtcA and NtcB boxes are colored in green and cyan, respectively. The numbers on the top indicate the positions of NtcA and NtcB relative to transcription start site. Sequence logos derived from the multiple-sequence alignment.

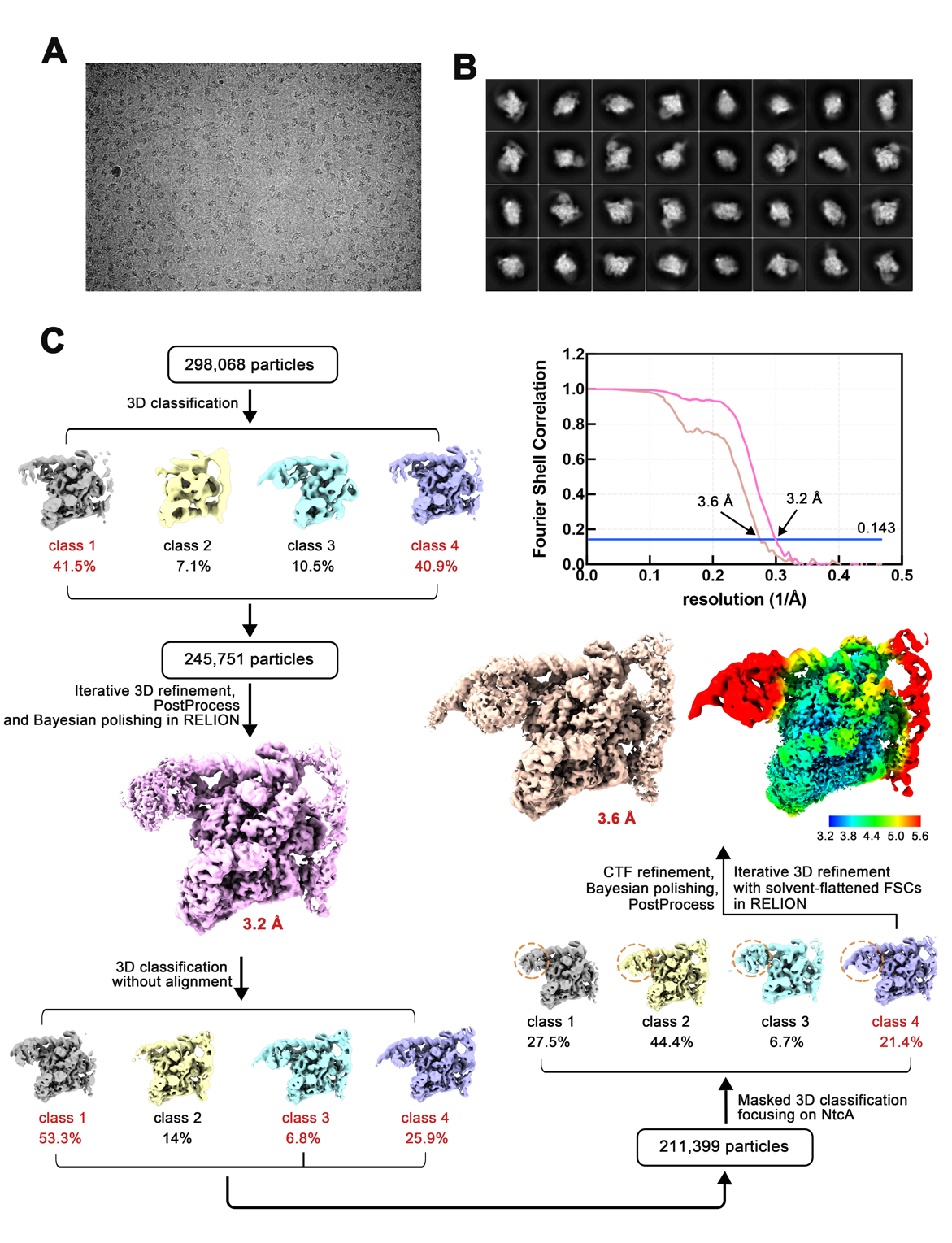

**fig. S2. Cryo-EM data processing of NtcA-TAC.** (**A**) A representative cryo-EM image of NtcA-TAC. (**B**) Representative 2D class averages of NtcA-TAC. (**C**) Flowchart for cryo-EM data processing and map reconstruction for NtcA-TAC.

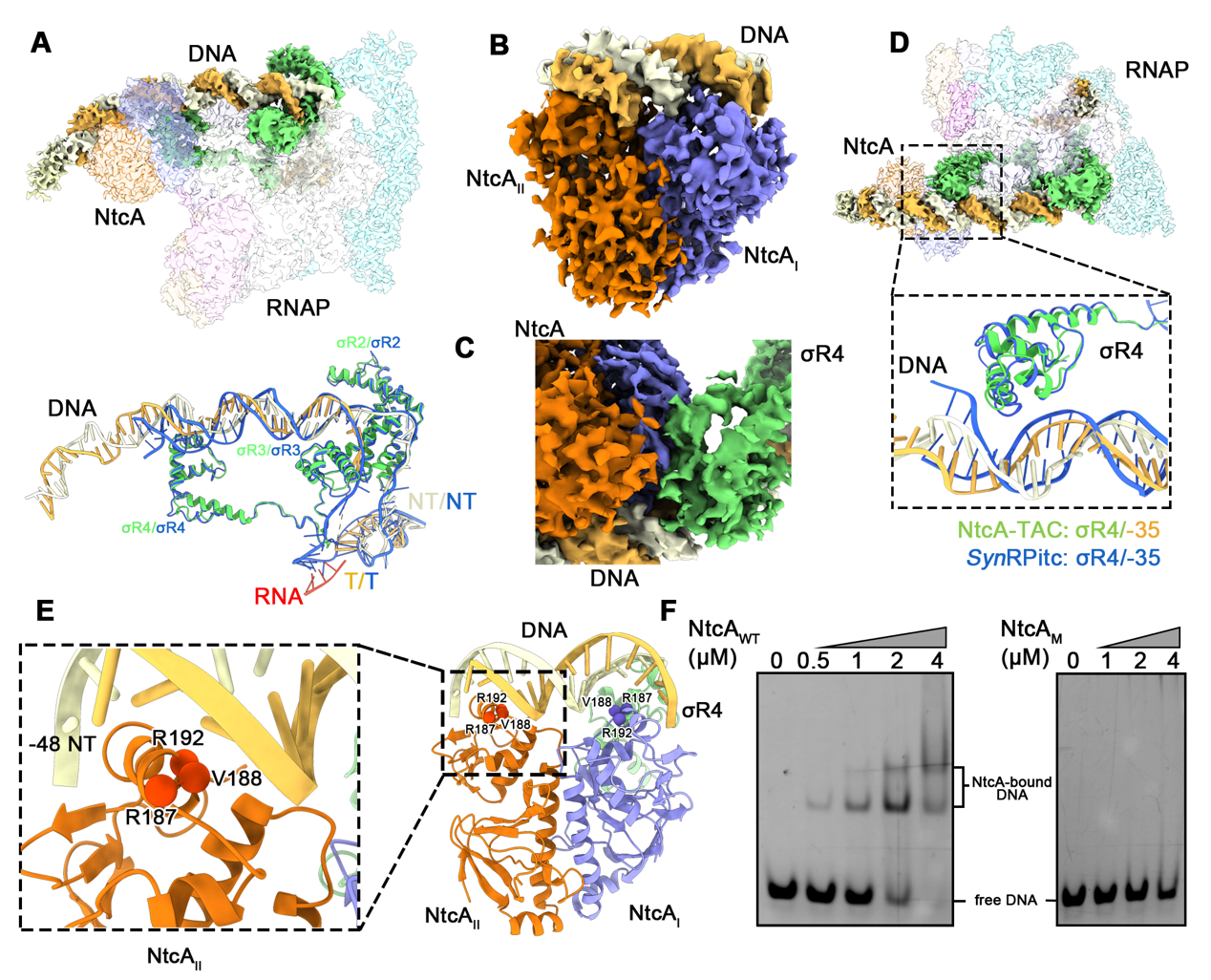

**fig. S3. The interaction pattern of NtcA binding to the promoter DNA and RNAP in NtcA-TAC.** (**A**) Superposition of promoter DNA between *Anabaena* NtcA-TAC and *Synechocystis* RPitc (*syn*RPitc, PDB: 8GZG). The color schemes are as follows: NT (nontemplate DNA) of NtcA-TAC, light yellow; T (template DNA) of NtcA-TAC, light orange; σ^A^ of NtcA-TAC, green; NT, T and σ^A^ of *syn*RPitc, blue; RNA of *syn*RPitc, red. The σR2, σR3, and σR4 represent the region 2, region 3 and region 4 of σ^A^, respectively. (**B**) Cryo-EM map of NtcA and its binding boxes. (**C**) Cryo-EM maps of NtcA-σR4. (**D**) Superposition of σR4-DNA between NtcA-TAC and *syn*RPitc. (**E**) The interactions between NtcA and the NtcA box of promoter DNA. The key residues of NtcA involved in binding to DNA are shown as spheres. (**F**) The EMSA results showing binding of wild-type (NtcA_WT_) or mutant (R187A/V188A/R192A; NtcA_M_) NtcA with a DNA fragment containing the NtcA box.

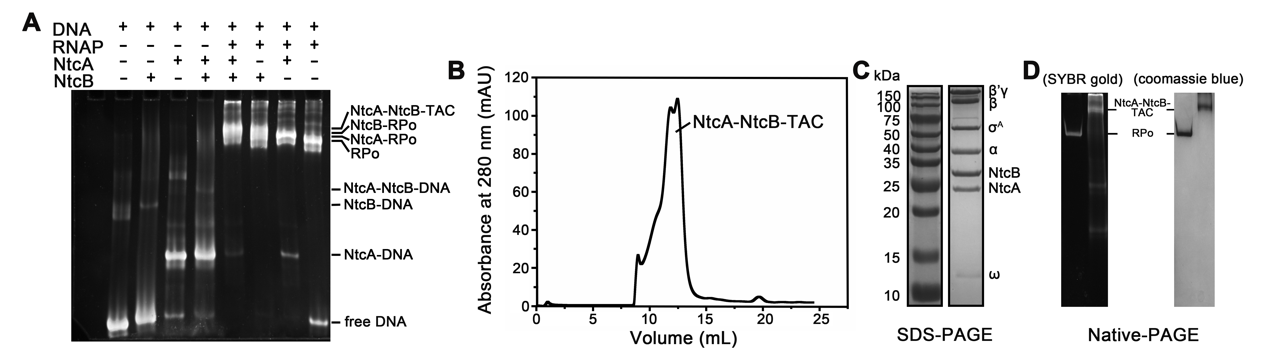
**fig. S4. In vitro reconstitution of NtcA-NtcB-TAC.** (**A**) Gel-shift result showing migration of protein-DNA complex reconstituted through indicated combinations of NtcA, NtcB, RNAP and the DNA in Fig. 1A. (**B**) The size-exclusion chromatography of NtcA-NtcB-TAC. (**C**) The SDS-PAGE analysis of the purified NtcA-NtcB-TAC complex. The protein components are labeled on the right of the gel. (**D**) The native-PAGE analysis of NtcA-NtcB-TAC stained with SYBR Gold dye (left) or Coomassie Brilliant Blue (right).

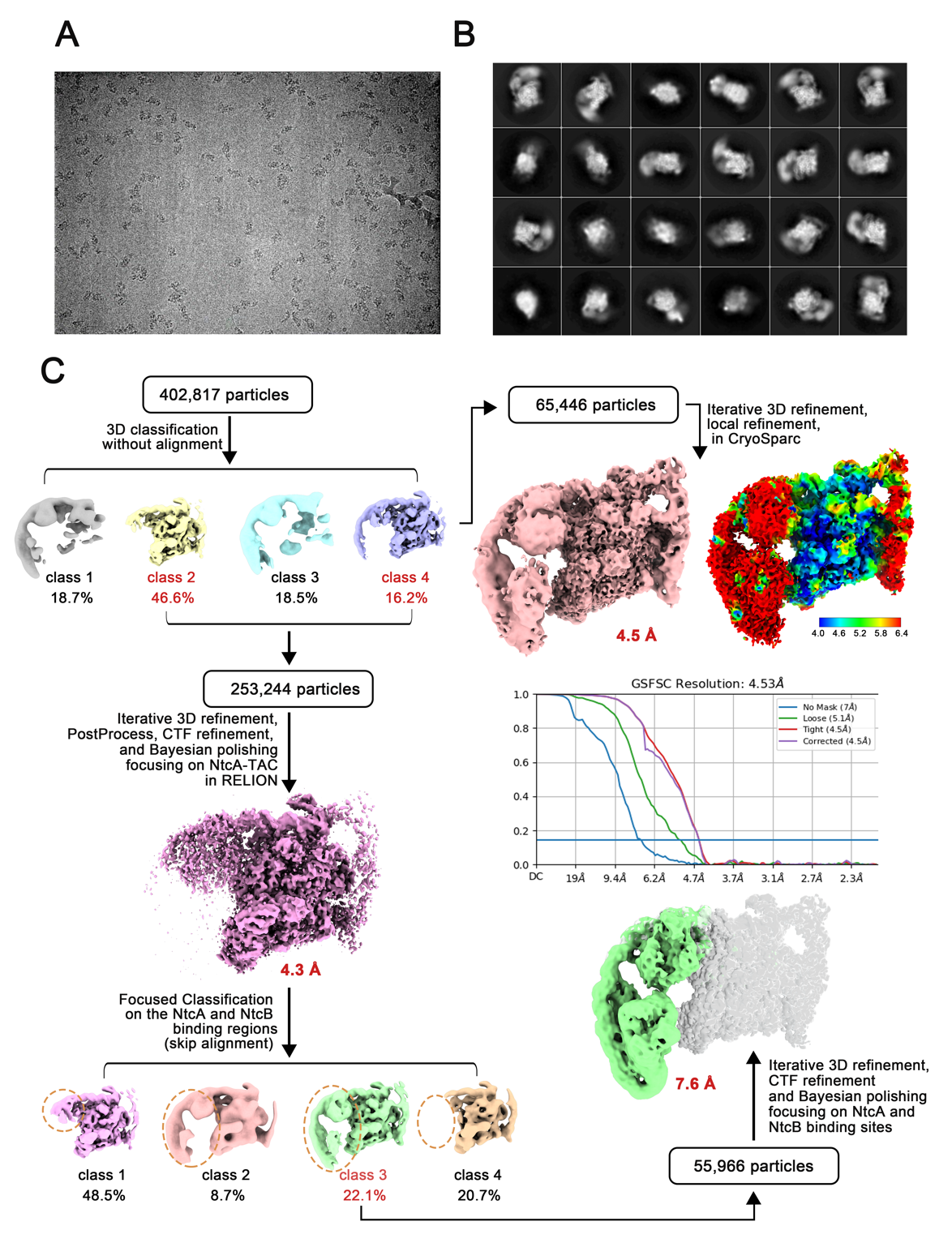

**fig. S5. Cryo-EM data processing of NtcA-NtcB-TAC.** (**A**) A representative cryo-EM image of NtcA-NtcB-TAC. (**B**) Representative 2D class averages of NtcA-NtcB-TAC. (**C**) Flowchart for cryo-EM data processing of NtcA-NtcB-TAC.

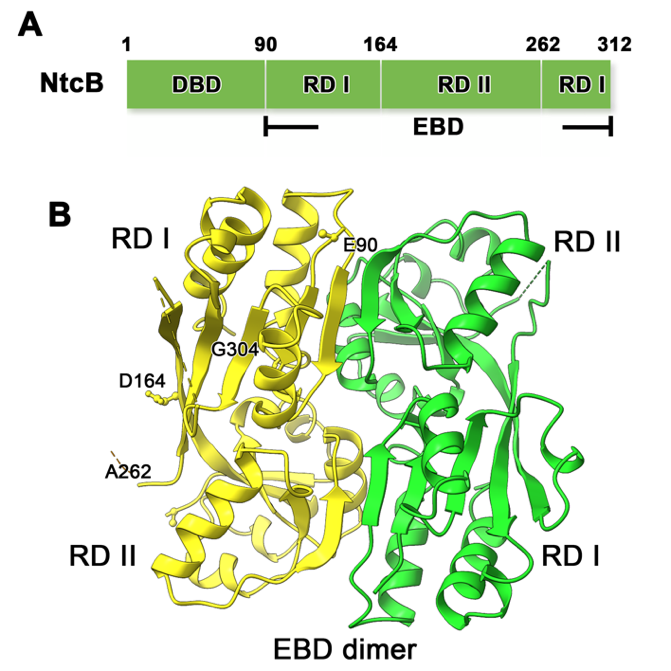

**fig. S6. The crystal structure of NtcB-EBD** (**A**) A schematic presentation of domain organization of NtcB. (**B**) Crystal structure of the NtcB-EBD dimer. The two subunits are colored yellow and green, respectively. The two subdomains RD I and RD II are labeled.

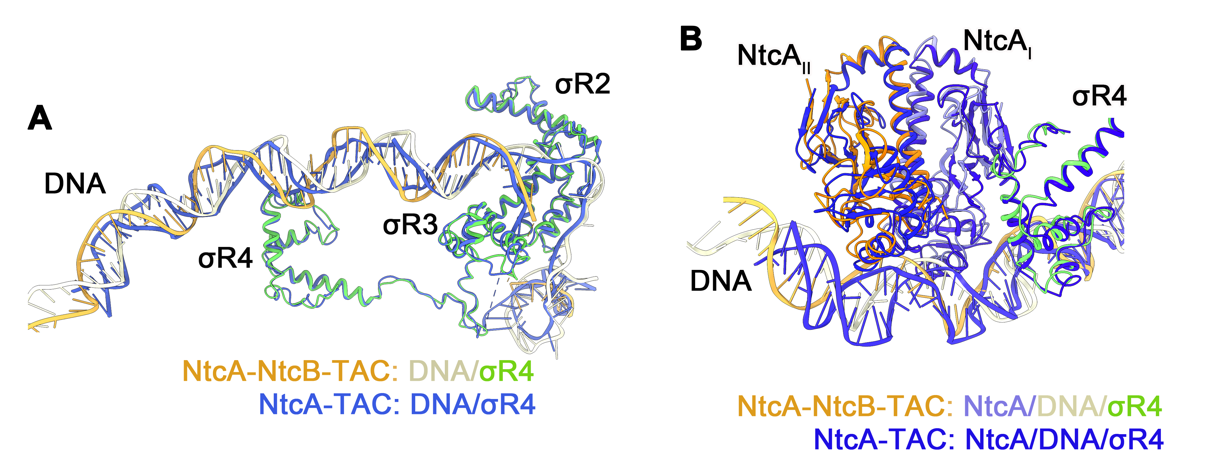

**fig. S7.** **Structural comparison of NtcA-TAC and NtcA-NtcB-TAC.** (**A**) Superposition of promoter DNA between NtcA-TAC and NtcA-NtcB-TAC. (**B**) Superposition of the NtcA-σR4-DNA between NtcA-TAC and NtcA-NtcB-TAC.

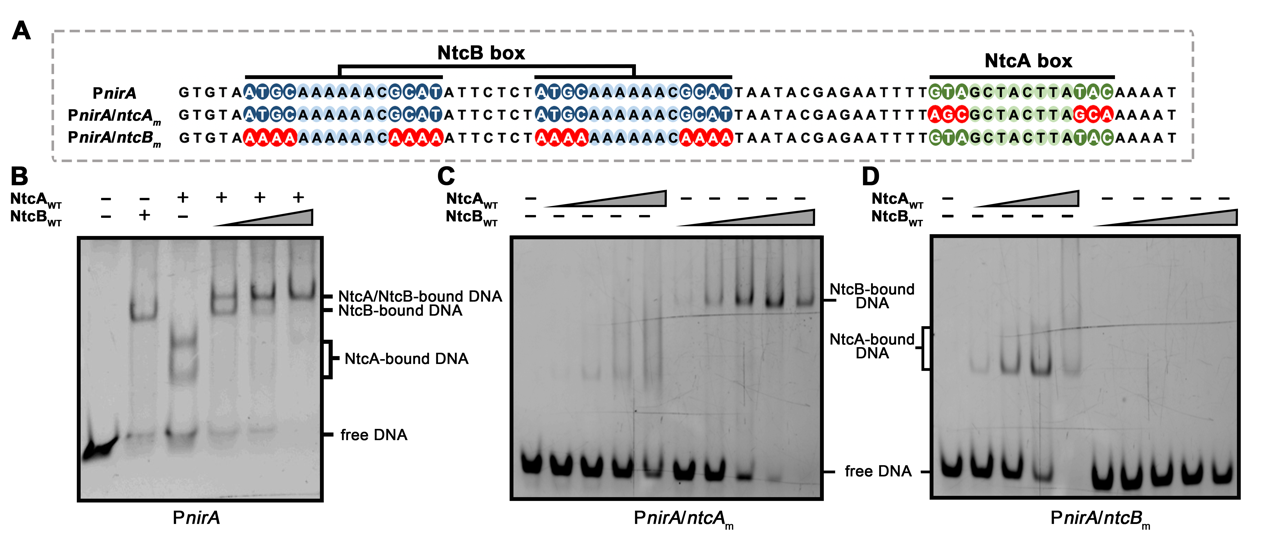
**fig. S8. NtcA and NtcB independently bind to their respective boxes.** (**A**) The sequences of 5'-Cy5 labeled dsDNA that are used in the EMSA assays. P*nirA*, 76-bp DNA containing wild type NtcA and NtcB boxes; P*nirA/ntcA_m_*, a P*nirA* derivative bearing mutations at the conserved palindromic nucleotides of NtcA box; P*nirA/ntcB_m_*, a P*nirA* derivative bearing mutations at the conserved palindromic nucleotides of NtcB box. The mutated nucleotides are colored in red. (**B**) NtcA and NtcB bind to P*nirA* independently. NtcA_WT_, wide-type NtcA protein; NtcB_WT_, wide-type NtcB protein. **(C)** Mutating the binding boxes of NtcA abolished NtcA binding activity but did not affect NtcB binding. **(D)** Mutating the binding boxes of NtcB abolished NtcB binding activity but did not affect NtcA binding.

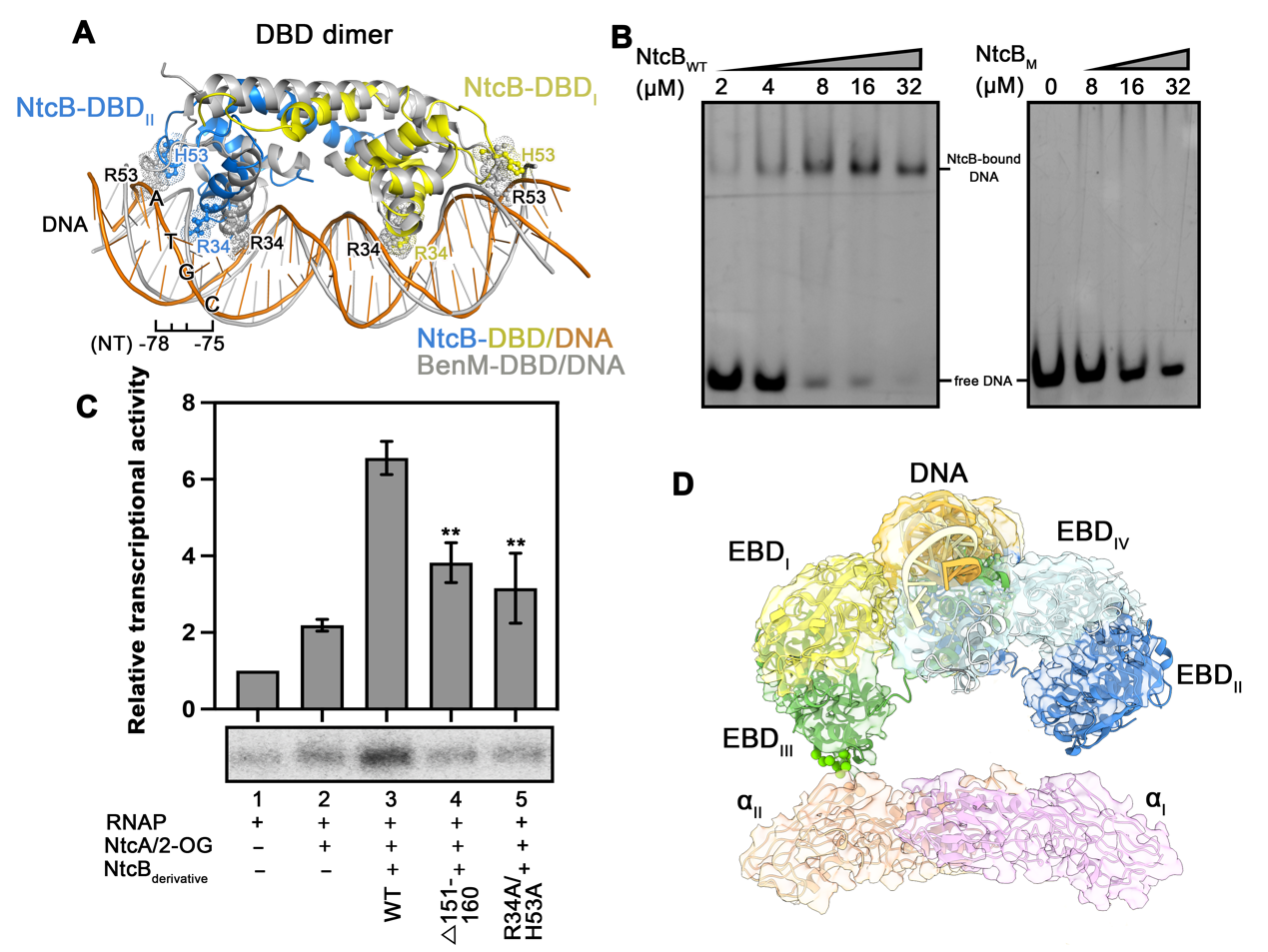

**fig. S9. The interactions of NtcB to promoter DNA and RNAP in NtcA-NtcB-TAC**. (**A**) Superposition of NtcB-DBD/DNA in NtcA-NtcB-TAC and the crystal structure of BenM-DBD/DNA complex (PDB:4IHS). The residues of NtcB (blue, yellow) and BenM (black) responsible for DNA motif recognition are labeled. (**B**) Alanine mutation of the DNA motif-recognition residues (R34A/H53A; NtcB_M_) of NtcB abolished the binding ability of NtcB to a DNA fragment containing NtcB boxes. (**C**) Mutations of the DNA motif-recognition or RNAP-contact residues of NtcB impairs the transcription activation activity of NtcB. Data are presented as mean ± S.E.M., n=3 biologically independent experiments. The lower panel shows the representative gel image. 2-OG, 2-oxoglutarate. △151-160, deletion of the contact patch (residues Leu151-Gly160) of NtcB. **P<0.01 indicates significant difference compared to lane 3. (**D**) Cryo-EM map showing the interface between RNAP and NtcB-αNTD.

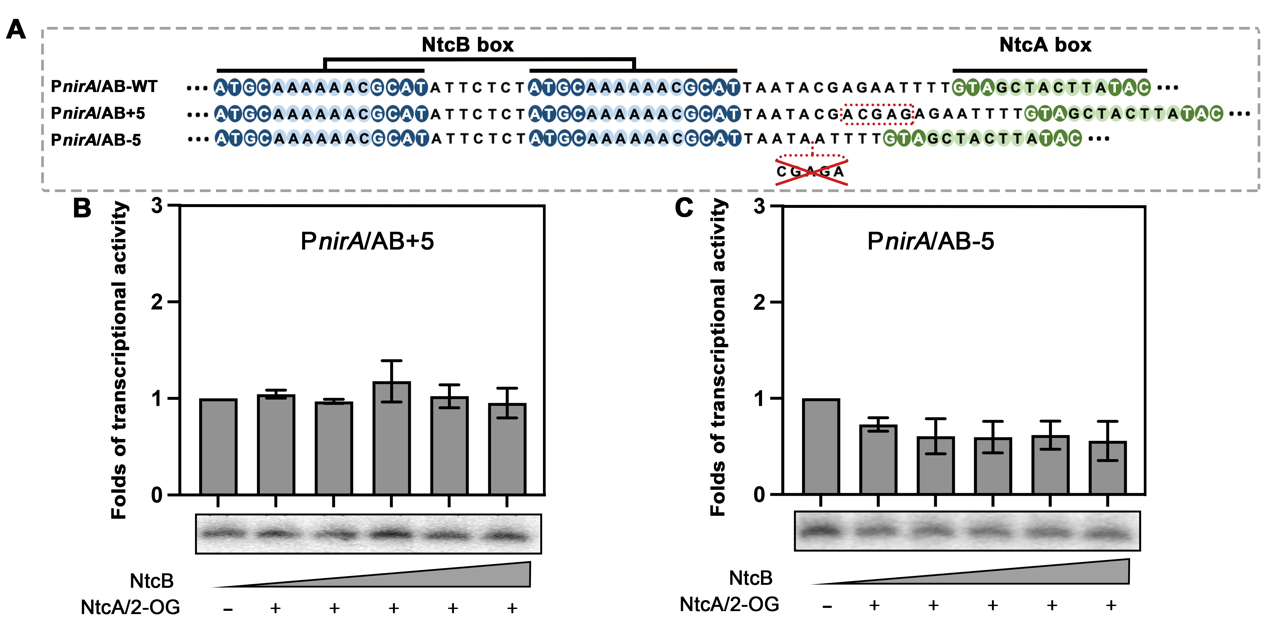
**fig. S10. Alteration of spacer between NtcA- and NtcB- boxes abolishes transcription activation activity of NtcB.** (**A**) The sequences of three promoter DNAs that are used in *in vitro* transcription assays. P*nirA*/AB–WT, P*nirA* containing wild-type spacer length between NtcA and NtcB boxes; P*nirA*/AB+5, a P*nirA* derivative containing insertion of the spacer between NtcA and NtcB boxes by 5 bp; P*nirA*/AB-5, a P*nirA* derivative containing deletion of the spacer between NtcA and NtcB boxes by 5 bp. The sequences of insertion or deletion of nucleotides are highlighted. (**B**) Insertion or (**C**) deletion of the spacer by 5 bp between NtcA and NtcB boxes impaired the transcription activation activity of NtcB. Data are presented as mean ± S.E.M., n= 3 biologically independent experiments. The lower panel shows the representative gel image.

**table S1.** The plasmid and DNA used in this study.

| **Oligonucleotides** | | | **Sources** |
| --- | --- | --- | --- |
| pET28a/*ntcA* | | | This study |
| pET28a/*ntcB* | | | This study |
| pET28a/*σ^A^* | | | This study |
| pET28a/*ntcB-EBD* | | | This study |
| pEASY/P-*nirA* | | | This study |
| pEASY/P-*nirA^AB+5^* | | | This study |
| pEASY/P-*nirA^AB-5^* | | | This study |
| PETDuet/*rpoA-rpoZ* | | | This study |
| PCDFDuet/*rpoB-rpoC1C2* | | | This study |
| **EMSA** | P*nirA*-F  (5′-FAM) | | GTGTAATGCAAAAAACGCATATTCTCTATGCAAAAAACGCATTAATACGAGAATTTTGTAGCTACTTATACAAAAT |
|  | P*nirA*-R | | AATTTTGTATAAGTAGCTACAAAATTCTCGTATTAATGCGTTTTTTGCATAGAGAATATGCGTTTTTTGCATTACAC |
|  | P*nirA*/*ntcA_m_*-F  (5′-FAM) | | GTGTAATGCAAAAAACGCATATTCTCTATGCAAAAAACGCATTAATACGAGAATTTTAGCGCTACTTAGCAAAAAT |
|  | P*nirA*/*ntcA_m_*-R | | ATTTTTGCTAAGTAGCGCTAAAATTCTCGTATTAATGCGTTTTTTGCATAGAGAATATGCGTTTTTTGCATTACAC |
|  | P*nirA*/*ntcB_m_*-F  (5′-FAM) | | GTGTAAAAAAAAAAACAAAAATTCTCTAAAAAAAAAACAAAATAATACGAGAATTTTGTAGCTACTTATACAAAAT |
|  | P*nirA*/*ntcB_m_*-R | | ATTTTGTATAAGTAGCTACAAAATTCTCGTATTATTTTGTTTTTTTTTTAGAGAATTTTTGTTTTTTTTTTTACAC |
| **DNA**  **scaffold** | | NT-DNA | GTTAAGTGTAATGCAAAAAACGCATATTCTCTATGCAAAAAACGCATTAATACGAGAATTTTGTAGCTACTTATACAAAATTCAGGAAAATTTTTCTGTATAATGGGAGCTGTCACGGATGCAGG |
|  |  | T-DNA | CCTGCATCCGTGAGTCGAGGGTAATAACAGAAAAATTTTCCTGAATTTTGTATAAGTAGCTACAAAATTCTCGTATTAATGCGTTTTTTGCATAGAGAATATGCGTTTTTTGCATTACACTTAAC |

**table S2.** Crystal parameters, data collection and structure refinement.

| Data collection | NtcB-EBD (PDB: 8H3Z) |
| --- | --- |
| Space group | *P2_1_2_1_2_1_* |
| Unit cell (Å, º) | 59.697 68.410 114.310 90.00 |
| Resolution range (Å) | 50.00-2.40 (2.49-2.40)^a^ |
| unique reflections | 34,980 (3,444) |
| I/σ(I) | 20.7 (2.8) |
| Rpim (%) | 5.4 (32.9) |
| CC_1/2_ (%) | 89.3 (78.7) |
| Completeness (%) | 99.0 (97.0) |
| Average redundancy | 8.3 (5.8) |
| Structure refinement |  |
| Resolution range (Å) | 50.00-2.40 |
| R-factor^b^/R-free^c^ (%) | 20.4/24.3 |
| Number of atoms |  |
| Proteins | 3,187 |
| Waters | 115 |
| RMS deviation from ideality |  |
| Bond lengths (Å) | 0.013 |
| Bond angles (º) | 1.576 |
| Average B factors (Å^2^) | 35.6 |
| Ramachandran statistics^d^ |  |
| Favored regions (%) | 98.23 |
| Allowed regions (%) | 1.77 |
| Outliers (%) | 0 |
| Validation |  |
| MolProbity score | 1.85 |
| Clashscore | 8.02 |
| Poor rotamers | 0.09 |

^a^The values in parentheses refer to statistics in the highest bin.

^b^R-factor =∑_h_|Fo(h)－Fc(h)|/∑_h_Fo(h), where |Fo| and |Fc| are the observed and calculated structure-factor amplitudes, respectively.

^c^R-free was calculated with 5% of the data excluded from the refinement.

^d^Categories were defined by Molprobity.

**table S3**. Cryo-EM parameters, data collection and refinement statistics

| Data collection and processing | NtcA-TAC  PDB: 8H40  EMD-34476 | NtcA-NtcB-TAC  PDB: 8H3V  EMD-34475 |
| --- | --- | --- |
| Magnification | 81,000 | 81,000 |
| Voltage (keV) | 300 | 300 |
| Electron exposure (e^–^/Å^2^) | 50 | 50 |
| Defocus range (μm) | -1.2 to -2.2 | -1.2 to -2.2 |
| Pixel size (Å) | 1.07 | 1.07 |
| Symmetry imposed | C1 | C1 |
| Initial particle images (no.) | 566,246 | 851,868 |
| Final particle images (no.) | 45,239 | 65,256 |
| Map resolution (Å) | 3.6 | 4.5 |
| FSC threshold | 0.143 | 0.143 |
| Map resolution range (Å) | 2.02~999 | 2.02~999 |
| **Refinement** |  |  |
| Real-space correlation coefficient | 0.88 | 0.77 |
| Initial model used (PDB code) | 8GZG | 8GZG |
| Map sharpening B factor (Å2) | -96.62 | -246.01 |
| **Model composition** |  |  |
| Non-hydrogen atoms | 34,919 | 46,335 |
| Protein residues | 4,130 | 5,326 |
| nucleotide | 124 | 221 |
| Waters | 0 | 0 |
| ligands | 0 | 0 |
| **RMS deviation from ideality** |  |  |
| Bond lengths (Å) | 0.073 | 0.053 |
| Bond angles (°) | 4.379 | 4.061 |
| **Validation** |  |  |
| MolProbity score | 3.91 | 4.57 |
| Clash score | 205.16 | 215.73 |
| Poor rotamers (%) | 0.85 | 5.63 |
| **Ramachandran statistics** |  |  |
| Favored regions (%) | 53.4 | 50.0 |
| Allowed regions (%) | 43.0 | 45.2 |
| Outliers (%) | 3.6 | 4.8 |
